# Drivers of plant-associated invertebrate community structure in West-European coastal dunes

**DOI:** 10.1101/2024.06.24.600350

**Authors:** Ruben Van De Walle, Maxime Dahirel, Ward Langeraert, Dries Benoit, Pieter Vantieghem, Martijn L. Vandegehuchte, François Massol, Dries Bonte

## Abstract

The organisation of species assemblages is affected by environmental factors acting at different spatial scales. To understand the drivers behind the community structure of invertebrates associated with marram grass -the dominant dune-building ecosystem engineer in European coastal dunes-, we set up a stratified sampling scheme in six biogeographic sectors along the North Sea. We tested to which degree local invertebrate species composition is affected by the spatial organisation of marram grass tussocks. We used distance-based RDA and a joint species distribution modelling approach to understand how species traits and their phylogeny contribute to invertebrate community composition. We show biogeography to be the most important driver, followed by species-specific responses to marram grass cover and vitality. Traits and phylogeny had a minor influence on the species distribution patterns. The residual species covariation suggests negative interactions between groups of specialist and generalist species. From an applied perspective, our research indicates that the biological value of nature-based solutions for the restoration and design of coastal dunes can be optimized by the design of a heterogeneous marram grass planting scheme and/or development.

## Introduction

Biodiversity is shaped by a complex interplay of biogeographic, regional, and local factors, each operating at a different spatial scale (Shmida and Wilson 1985, Heino et al. 2015). Biogeographical factors, acting over geological time scales, influence colonisation-extinction dynamics, speciation, and species distribution along climatic gradients, for example with changing latitude and elevation (Whittaker et al. 2017). Regional diversity of terrestrial species is shaped by environmental heterogeneity, including soil type, topography and connectivity of local communities through dispersal (Leibold et al. 2004). At a local scale, species coexistence and community composition is further shaped by antagonistic, mutualistic and competitive interactions (McGill 2010, Patrick and Swan 2011, De Araujo et al. 2023). Closely related species with similar interaction traits may actively avoid each other at local scales to reduce competition for resources, yet they may still exhibit positive associations at regional scales due to shared habitat characteristics (Mayfield and Levine 2010, Gerhold et al. 2015).

Herbivore abundances and traits shape community dynamics through interactions with their host (Polis and Strong 1996). Plant species, particularly through their nutritional value and defence mechanisms, directly affect herbivore community composition (Mertens et al. 2021). Plant-associated invertebrate species may compete with one another, either directly for food resources, or indirectly by means of the host plant’s defence mechanisms. Alterations in plant defences in response to herbivory can have cascading effects through the community (Bezemer and van Dam 2005, Leimu and Koricheva 2006). While interactions primarily occur within compartments of the system, they can also extend across compartments, such as above-belowground interactions in terrestrial systems (Wardle et al. 2004).

The structural organisation of the vegetation, often referred to as structural complexity, directly affects the realised interaction strengths among competitors and between prey and their predators (Ritchie and Olff 1999). It also shapes non-trophic niches, e.g., through its impact on the microclimate (Rietkerk et al. 2004). Structural complexity emerges vertically within the vegetation, but also horizontally from the cover and patchiness of the different vegetation units. Island-mainland or metacommunity processes that are typically associated with the organisation of biodiversity at regional scales (Leibold et al. 2022) also act at local scales. In a managed meadow system, for instance, the diversity and abundance of arthropods was shown to directly depend on the size of local unmanaged vegetation patches and their distance to larger unmanaged area (e.g., the mainland source) of the system (Harvey and MacDougall 2014). Heterogeneous habitats therefore have the ability to support more species (Stein et al. 2014), as long as increasing heterogeneity does not come at the cost of a diminished area per habitat type (i.e. the area-heterogeneity trade-off; Allouche et al. 2012).

A complementary focus on species traits and individual species identity allows for a more mechanistic understanding of the processes shaping species communities (McGill et al. 2006, Wong et al. 2019). A trait commonly associated with the environmental filtering of arthropods is body size (Wong et al. 2019), because it is linked with important processes such as dispersal, predation and thermoregulation (Gravel et al. 2013, Hillaert et al. 2018, Pincebourde et al. 2021, Logghe et al. 2023). Species can also be classified according to their biotic interactions in so-called functional groups, which can be used to study interactions among functional groups rather than between actual species (Wong et al. 2019). Phylogenetically related species are more likely to have similar traits and occurrence patterns (Futuyma and Kirkpatrick 2018). Hence, incorporating phylogenetic relationships to account for similarity in unmeasured traits (Grafen 1989, Futuyma and Kirkpatrick 2018) can further improve our understanding of the processes shaping community assembly.

Coastal dunes in the Western Palearctic, dominated by European marram grass (*Calamagrostis arenaria*), exhibit low plant species diversity. Marram grass, a key ecosystem engineer, stabilises sand and forms monocultures along sandy coasts (Bonte et al. 2021). Its abundance is influenced by local sand dynamics, with patchy spatial clustering (Bonte et al. 2021). While this horizontal structural complexity shows large variation, vertical structural complexity is mainly determined by the extent of litter accumulation at the ground level. With waning sand dynamics, marram grass loses its vigour, most likely because root-attacking soil biota are largely absent from wind-blown beach sand but accumulate in the root zone in the absence of aeolian sand deposition (Huiskens 1979, van der Putten 1990).

Human interventions, such as planting or plant removal, can alter the horizontal structure of marram vegetation, but biogeomorphological processes largely drive it. Invertebrate species richness within marram dunes is limited due to environmental stressors, but most species positively associate with marram grass for feeding and shelter (McLachlan 1991, Maes and Bonte 2006). Biogeographical clustering of invertebrate communities is expected due to climatic gradients, dune geological history and limited connectivity. The geological history of coastal dunes from France, Belgium, the Netherlands, and Great Britain is strongly linked to geological connectivity with older sandy and/or limestone regions, the presence of calcareous or acidic sediments and dune development (Bonte et al. 2003a). At the meso-scale, metacommunity processes related to the landscape context or local management may be important as well. Environmental filters on invertebrate community composition are therefore expected to act at the local, regional and biogeographical scales. The relative structural simplicity of marram grass vegetation, varying mainly in only two dimensions, the well-studied drivers of this structural heterogeneity, and its tractable invertebrate community make marram vegetations ideally suited to test hypotheses regarding the role of spatial heterogeneity in driving community composition.

Species traits, such as body size, and phylogenetic relationships, summarising unmeasured traits, are likely to affect species responses to environmental variables, and thus shape community assembly. Understanding the relative importance of biogeographical, regional, and local factors in shaping invertebrate community structure in marram grass dunes involves considering environmental predictors, species co-occurrences, and regional biogeographic variation. Key research questions include the extent of variation in species occurrence due to environmental variables versus random processes, the influence of species traits and phylogenetic relationships on species responses to environmental variables, and the structure of species co-occurrence networks.

## Material & Methods

### Study area & design

We studied coastal dunes in Belgium, the Netherlands, the south of the United Kingdom and the north of France, predominantly along the Channel and the southern part of the North Sea. The studied dune systems can be divided into six distinct biogeographic sectors, which differ in soil and vegetation characteristics because of their geological history and climate (Bonte et al. 2003a) and may therefore host different invertebrate species pools. These six areas are: North Devon and Norfolk coast in the United Kingdom (the main regions in Southern UK with well-developed coastal dunes, rather than rocky shores), the Boulonnais region in France from Camiers until Dunkerque, the Flemish dune region from Dunkerque until Knokke, the Renodunaal region from Cadzand until Bergen aan Zee, and the Wadden district in the Netherlands from Bergen aan Zee to Texel (Fig. 1). Although all sampled blond dunes are somehow calcareous, the latter two regions are substantially more acidic relative to the other studies locations. Boulonnais and Flemish dune sands are especially rich in lime. Insularity separates the (calcareous) UK sites from the other ones. We give an overview of the average climatological conditions of these six regions in Appendix 1.

**Figure 1.**
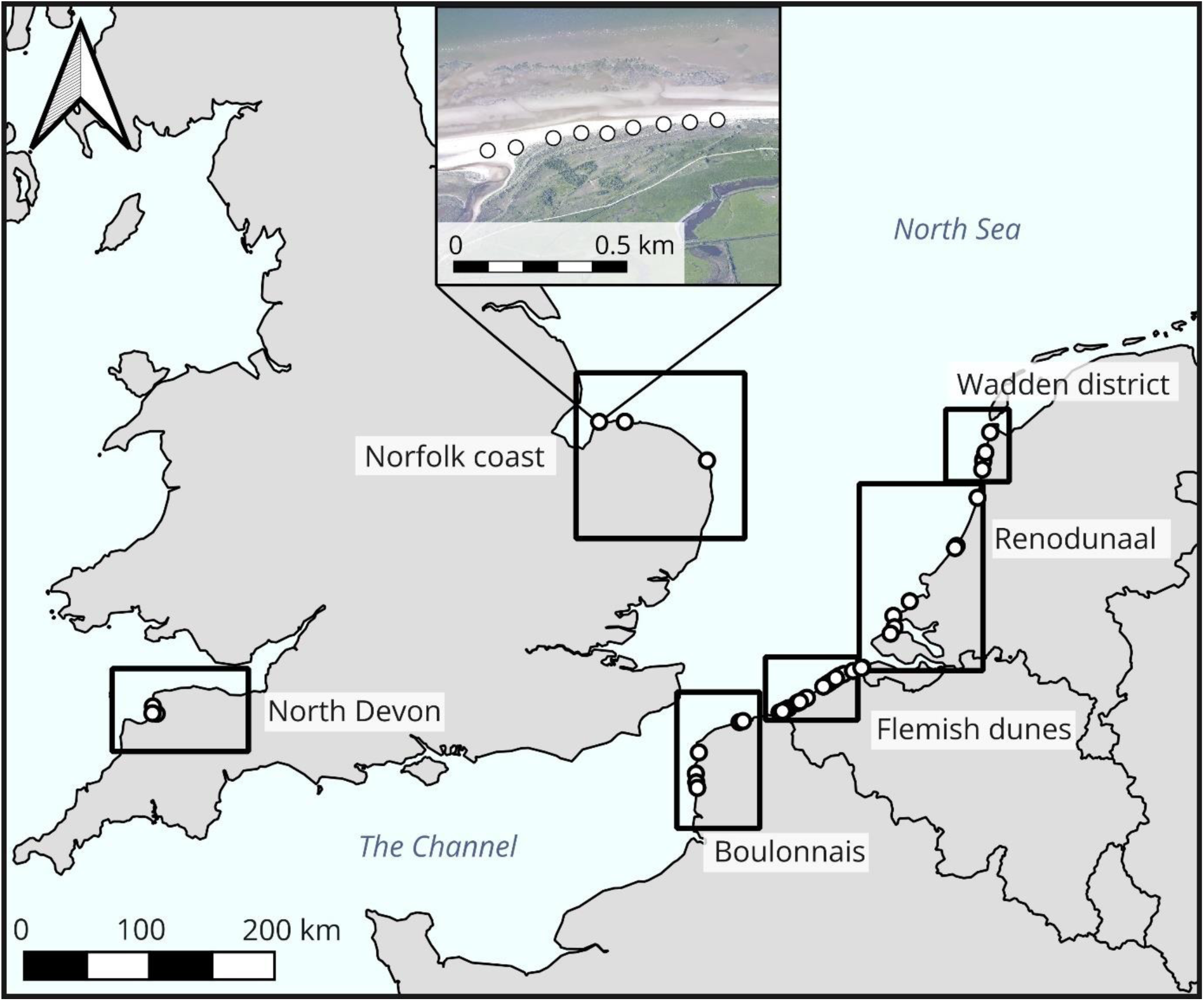
Geographical scope of the study. Dots denote the centroid locations of individual transects, grouped by biogeographic sectors. © EuroGeographics for the administrative boundaries. Inset: example of sampling unit locations within a given transect (Holme, UK); aerial photograph from the UK Environment Agency, under Open Government Licence.

In each biogeographic sector, transects parallel to the coastline and within 100m from the driftline were drawn in locations with sandy coasts and marram-dominated, blond dunes (Natura 2000 habitat 2120, CORINE biotope 16.21). Transect lengths were variable and chosen on site to incorporate the available variation in marram grass cover (average length: 1212 ± 786m). Given the strong fragmentation of the dune areas into separated entities due to recent urbanisation (usually a reserve in between coastal villages or cities), one entity was sampled by a single transect. The number of samples along each transect depended on the length of the transect, with individual samples separated by at least 20 m. Each sample was centred on a marram grass (*Calamagrostis arenaria* (L.) Roth) tussock surrounded by only marram grass vegetation and bare sand (e.g., no shrubs, trees, or large quantities of other species) in a radius of 5 m. Sampling units were chosen to maximise the variation in (1) vitality of the central marram grass tussock and (2) the spatial configuration of the surrounding marram grass cover (see Explanatory variables below). A total of 638 tussocks were sampled across all regions during the summers of 2017-2019.

### Invertebrate sampling

At each sampled marram grass tussock, aboveground invertebrates were sampled by sweep netting in and above the tussock for 15 seconds. Afterwards, ground-dwelling invertebrates were collected manually at the base of the tussock for 5 minutes. Sampling was only performed on relatively sunny days so flying insects would be active. All specimens were stored in 70 % ethanol, before being counted and identified using a stereomicroscope. Altogether, 15 726 individuals from 632 taxonomic units were identified, among which 434 taxa to species level, 96 to genus level and 102 to family level or higher. The used ID keys and trait sources are supplied in Appendix 2.

### Explanatory variables

We considered four sources of explanatory variables in our analyses: (1) regional characteristics (spanning multiple transects) (2) local environmental variables (at sampling location level), (3) species traits, and (4) phylogenetic relationships between species.

#### Regional environmental variable

The name of the biogeographic sector (as described above) was included as the only large-scale environmental variable.

#### Local environmental variables

For each sample, we first assessed the vitality (V) of the focal marram grass tussock. Vitality was initially noted as a 5-category score (from 0 to 4) based on the estimated proportion of green visible in a photograph of the central marram grass tussock. However, low vitality tussocks were very rare in the dataset (1 zero and 22 ones out of 588 samples); the 0 and 1 categories were therefore grouped together in subsequent analyses. We then quantified the proportional marram grass cover (P%) and spatial clustering (e.g., correlation) of marram grass surrounding the central tussock, using high-resolution vegetation maps of the coastal dune areas (Bonte et al. 2021). Both calculations were done within a 50 m radius circle around each focal marram grass tussock. We used Moran’s *I* as a measure of spatial autocorrelation (Moran 1950, Bivand and Wong 2018), with negative values indicating an increasingly regular configuration, zero a random configuration and positive values an increasingly clustered configuration. Moran’s *I* values were calculated using the “moran.test” function from the *spdep* R package (Bivand and Wong 2018). Marram grass naturally grows in clusters (Bonte et al. 2021), so Moran’s *I* values in the analysed dataset were high, ranging from 0.75 to 0.98.

#### Species traits

For each species, we included the average body length (in millimetres) and the functional group at the adult stage (detritivore, herbivore, predator or omnivore) as trait variables. Data were obtained from direct measurements and the literature (Van De Walle et al. 2023, Logghe et al. 2023; see Appendix 1).

#### Phylogenetic relationships

We used the R package *rotl* (Michonneau et al. 2016) to query the Open Tree Of Life (OpenTreeOfLife et al. 2019) for a phylogenetic tree connecting all 50 species (Fig. 2). The resulting tree does not contain branch length information, which is needed for analyses; we used Grafen (1989)’s method as implemented in the R package *ape* (Paradis 2019) to assign branch lengths.

**Figure 2.**
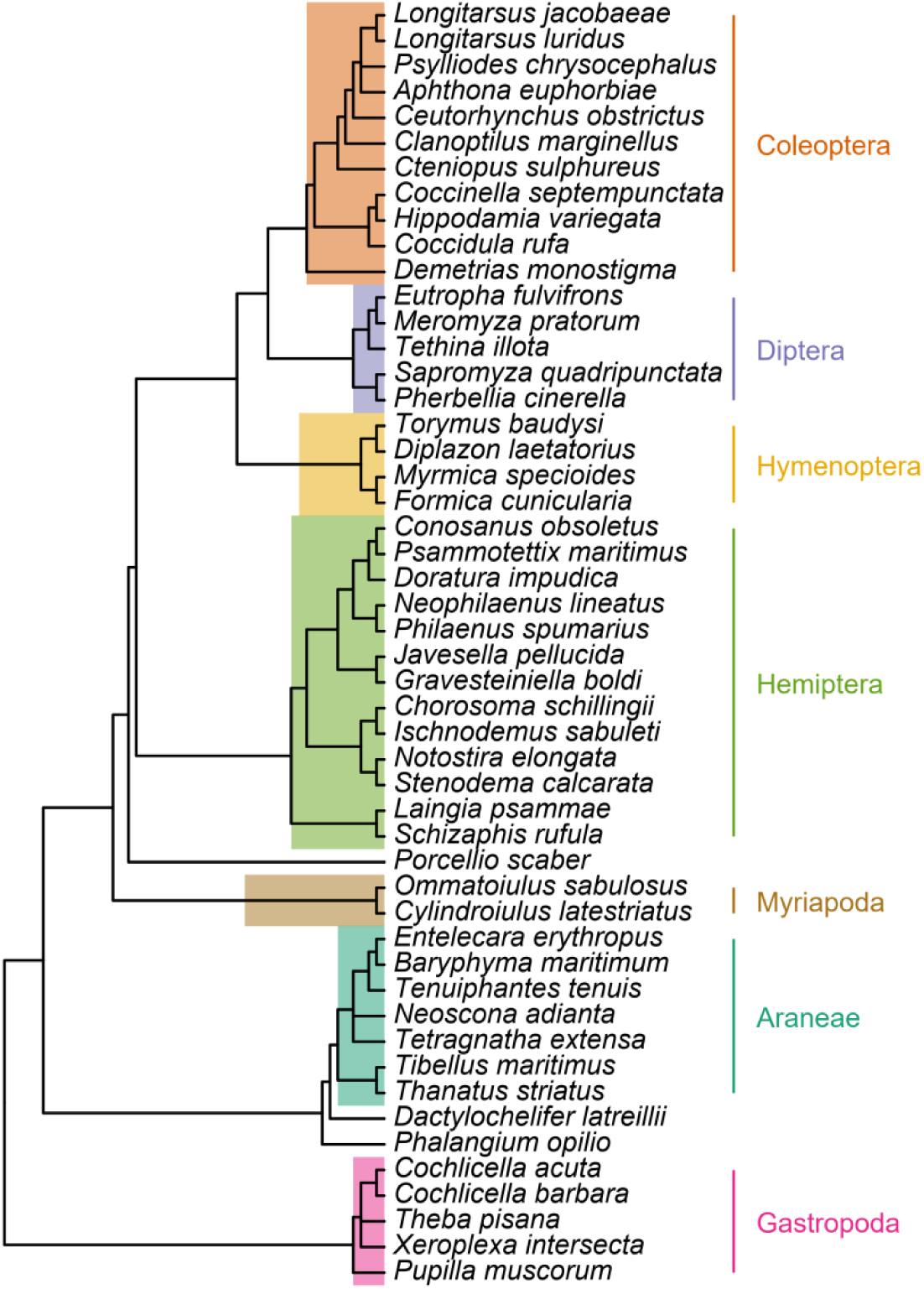
Phylogenetic tree linking the 50 species retained for analyses, annotated with major clade names, obtained from the Open Tree of Life (see text for details).

### Statistical analyses

Data were analysed using R (version 4.3.3, R Core Team 2024).

Initial data exploration revealed that species abundance data contained many zeros (i.e. most species were very rare), even after combining the abundances of both capture methods into one single count per species per sample. We applied and compared two different approaches to understand how invertebrate taxa are correlated with the presumed drivers of the community structure.

First, we used an abundance-based Redundancy Analysis (RDA) to identify the putative most important environmental variables (marram grass cover, Moran’s I, vitality, biogeographic sector) explaining variation in community composition. We added to the analysis the sample latitudes and longitudes to visualise the ordering according to the biogeographic sectors. The distance-based RDA used all species with > 15 individuals in the dataset, with 583 remaining samples. We subsequently applied a PERMANOVA analysis based on a Bray-Curtis distance matrix to quantify the variation explained by the included environmental factors on the full assemblage structure. Analyses were done using the RDA and Adonis2 functions as available in the *vegan* R package (Oksanen et al. 2022. We used a complex design with nested factors (order from transects in regions, local marram variables within transects) and follow a sequential SS (typeI) to ensure each term is fitted after the previous one. In such a way, PERMANOVA mimics to the best the random design implemented in the JSDM. For local variables, we chose to fit vitality before the spatial variables (but alternative sequences provided the same results). Permutations were run within the transect-“stratum”.

Second, to take full advantage of the complexity of the species co-occurrence by considering the single species responses, their traits and phylogeny in concert (Ovaskainen et al. 2017), we modelled the relationships between environmental variables, species characteristics including phylogeny and the occurrence of all target species simultaneously using Joint Species Distribution Modelling (JSDM).

More specifically, we used the Hierarchical Modelling of Species Communities framework (HMSC) as implemented in the *Hmsc* R package (Ovaskainen and Abrego 2020, Tikhonov et al. 2020a, b, Poggiato et al. 2021). To be able to draw reliable inference from the data, we focused on modelling occurrence only (absence = 0, presence = 1) rather than abundance, and kept only the most common species (Ovaskainen and Abrego 2020). After excluding samples with taxa not identified up to species level, the 50 most common species that occurred in at least 20 samples were selected. These 50 species covered two different phyla (Arthropoda and Mollusca with 45 and 5 species respectively, Fig. 2). This filtering reduced the number of marram grass tussocks used in analyses from the initial 638 to 588, within 44 transects.

In practice, HMSC models are Bayesian multi-response generalized linear mixed models (one species = one response), with the distribution and link function depending on the type of species data. As we analysed occurrence data, our model assumed responses were Bernoulli-distributed (presence/absence at the sample/tussock level) and used a probit link function. For a detailed overview of the internal structure of a HMSC model, including the articulation between fixed and random effects, or how traits and phylogeny influence niche parameters, see (Ovaskainen et al. 2017, Ovaskainen and Abrego 2020).

#### Environmental variables (fixed effects)

To account for regional differences in climate and soil or differences in species distribution originating from historical metacommunity processes, the model included the name of biogeographic sector as a factor. To account for local conditions, the model included (i) an effect of tussock vitality (V), implemented as a categorical factor, (ii) a continuous effect of spatial configuration (Moran’s *I*), and (iii) a continuous effect of marram grass cover (P%). To allow the latter to be non-linear (i.e. modelling optimal intermediate cover), it was included as a combination of linear and quadratic terms. We did not implement interactions between environmental variables, and neither included explicit spatial effects, since coordinates align with the biogeographic sector already considered as categorical factors and would lead to an overfitting and the subsequent convergence problems.

#### Species characteristics (fixed effects)

Both species body size and functional groups were included as covariates potentially explaining between-species variation in niches (i.e. in the effect of the environmental variables). This is implemented in practice as an internal linear sub-model explaining between-species variation in environmental niche coefficients (see previous paragraph) using trait values. The phylogenetic tree was also added (as a phylogenetic correlation matrix) to account for potential unmeasured and phylogenetically conserved traits that may explain species niches. This allows to estimate a phylogenetic signal parameter *ρ*, which is the proportion of the between-species covariance in responses to the environmental variables that is explained by their phylogenetic correlation.

#### Species “residual” association patterns (random effects)

Finally, we implemented sampling unit-level random effects, correlated among species, to account for species co-occurrence patterns not explained by environment, traits or phylogeny. As the number of correlation parameters becomes rapidly unwieldy as the number of species increases, JSDMs - including *HSMC* - use approaches based on latent factors (Ovaskainen and Abrego 2020). We initially attempted to use either spatially explicit random effects (Tikhonov et al. 2020b) or hierarchical random effects (sampling units nested in transects) to account for the spatial distribution of our sampling units and its potential effect on species co-occurrences. However, the model consistently failed to converge in both cases, with diagnostic values for latent factor parameters staying stuck even as chain length or thinning rate were increased, and priors altered from defaults (see below for diagnostics). We therefore implemented species co-occurrence as sampling unit-level non-spatial random effects. We evaluated post-hoc whether there could be spatial patterns by extracting the posterior values for the site latent factors, generating correlograms from each posterior sample (using the *ncf* package, Bjornstad 2022), and using the resulting 95 % credible intervals to determine if observed patterns could be explained from spatial structure.

All continuous environmental and trait variables were centred and scaled to unit 1 SD before inclusion in the model. This enables us first to be able to interpret linear and quadratic terms independently of each other, and second, to directly compare the effects of different variables on standardised units (Schielzeth 2010)

We kept the default priors (as described and justified in Ovaskainen and Abrego 2020) in most cases, with one exception: in the set of five parameters used to define the prior variance of the species loadings, and through that the residual co-occurrence matrix, priors for the parameters (a_1_, a_2_) were changed from their default of (50, 50) to (60, 60). This has the effect of adding more shrinkage to the species co-occurrence matrix compared to the default, making predicted association networks simpler and reducing overfitting (Ovaskainen and Abrego 2020). This enabled the model to converge in the first place when random effects were included, when it had difficulties doing so with default priors. This may be because our dataset is composed of species absent from most samples, even after the removal of most rare taxa (see Results), meaning that the data potentially contained limited information about co-occurrence.

We sampled the posterior distribution with four Markov Chain Monte Carlo (MCMC) chains. Each chain ran for 45000 iterations, with a burn-in period of 15000 iterations and a thinning factor of 30. This resulted in 1000 retained posterior samples per chain. We converted model outputs from the internal *HMSC* format to the format used in the *posterior* R package (Bürkner et al. 2023) to diagnose and summarise posteriors, with the format from the *coda* R package (Plummer et al. 2006) used as an intermediate step. We checked chain convergence using the updated version of the potential scale reduction factor (*R̂*) described in (Vehtari et al. 2021) and all *R̂* values were below 1.01 (Supplementary Fig. S5). We additionally checked and confirmed that both tail and bulk effective sample sizes as defined in Vehtari et al. (2021) were satisfactory (each > 400 for all parameters, and in most cases > 1000; Appendix 3).

We evaluated the model based on its out-of-sample predictive performance approximated through 10-fold cross-validation, using two metrics each calculated on a species per species basis. First, Tjur’s *D* (Tjur 2009) is a discrimination coefficient, which equals 1 if the model perfectly discriminates between sites where a species is present *vs* sites where it is absent, and 0 if it classifies sites at random. *D* can be seen as analogous to an R² statistic (Tjur 2009). Second and as a contrasting metric, we used the area under the precision-recall curve (AUC-PR, (Sofaer et al. 2019, Poisot 2023)). Contrary to Tjur’s *D* and to the often-used area under the receiver-operator curve (AUC-ROC), AUC-PR does not use true negatives in its calculation. This makes it relevant in joint species distribution models where most species are absent from most sites; in these contexts, even a poor model might reach high accuracy by being accidentally right about many absences. AUC-PR ranges from 0 to 1, with the expected value if the model classifies at random being variable and equal to the observed prevalence of the focal species.

When posterior summaries are given, they are given as “mean [2.5%, 97.5% quantiles]”.

## Results

### Multivariate analysis based on species abundances

The RDA shows a clear clustering of the samples according to the biogeographic sector, and their association with the continuous explanatory factors (Fig. 3). The first ordination axis explains 20% of the variation and is associated with longitude (separation of UK samples), as well as with local factors (marram cover and spatial clustering). These local factors are also correlated with the second axis (predictors explaining 12% of the variation). Factor covariation along both axis is evidently due to the nested sampling design that ensured we covered the same range of marram cover, clustering and vitality per transect and biogeographic sector. PERMANOVA analysis demonstrated the overall importance of biogeography as explaining variable of the community structure (R²=12%; Pseudo-F=18.35; *df*=5, *P*<0.001), but also the strong transect-dependent variation within the sectors (R²=18%; Pseudo-F=3.88; *df*=38, *P*<0.001). Additional within-transect variation was explained by the vitality (R²=7 %, Pseudo-F=1.29, *df*=43, *P*<0.001), cover (R²=7 %, *Pseudo-F*=1.24, *df*=44, *P*<0.001) and clustering (R²=6 %, Pseudo-F=1.13, *df*=43, *P*=0.003) of the marram grass tussocks.

**Figure 3.**
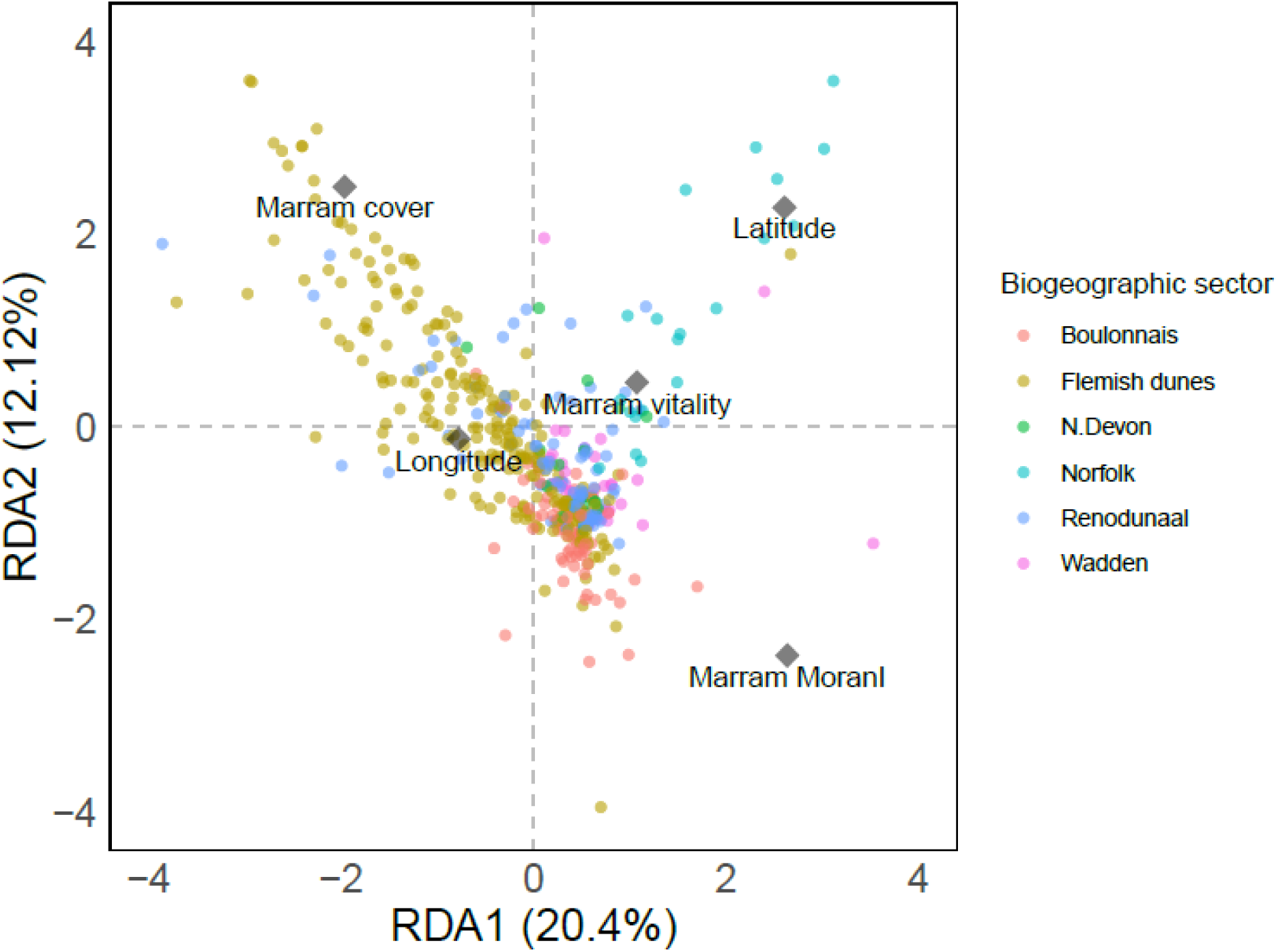
RDA-biplot with indication of the biogeographic sectors, and the steering environmental variables that influence the local invertebrate community structure.

### Species occurrence and JSDM model performance

Within the set of 50 “common” species kept in the JSDM analysis, the most prevalent species were *Theba pisana* (O. F. Müller, 1774), which was present in 36.2 % of the 588 samples, *Neophilaenus lineatus* (Linnaeus, 1758) (25.9%), *Demetrias monostigma* (Samouelle, 1819) (25.2%), *Meromyza pratorum* (Meigen, 1830) (21.6 %), and *Tibellus maritimus* (Menge, 1875) (20.2 %). The average species was present in 9.5 % of the samples (predicted value after 10-fold cross-validation: 9.4 % [9.1, 9.7 %]). The estimated out-of-sample predictive performance of our model, as measured after 10-fold cross-validation, was highly variable between species. Tjur’s *D* ranged from −0.00 [-0.01, 0.02] (*Conosanus obsoletus*) to 0.32 [0.29, 0.36] (*Meromyza pratorum*), with an average of 0.08 [0.08, 0.09] (Fig. S4-1). *D* values were positively correlated with species occurrence (*r* = 0.56 [0.50, 0.61]). AUC-PR values showed a similar spread, although the exact rankings between the top performing species were slightly different between the two metrics (Fig. S4-1). Using log odds ratios to control for the effect of observed occurrence on expected AUC-PR values, the performance of the model relative to random expectation is similar across most species (Fig. S4-2). Both Tjur’s *D* and AUC-PR show that out of sample, the model fails to predict occurrences better than the random expectation for four species: *Conosanus obsoletus*, *Tetragnatha extensa*, *Thanatus striatus* and *Torymus baudysi* (Fig. S4- 1).

### Variance partitioning from JSDM

For most of the species, variation explained by the model can be attributed to the biogeographic sector (Fig. 4, mean across species: 62.4 %, SD across species: 12.6 %). The local environmental variables together explained on average 24.3 % (SD: 9.3 %), with proportional cover of marram grass explaining 12.2 % (SD: 7.2 %), vitality of the marram grass 7.2% (SD: 4.7 %), and the spatial configuration 4.8 % (SD 4.9 %). The “residual” variation captured in the sampling unit-level random effect accounted for the remaining 13.3 % (SD: 9.4 %).

**Figure 4.**
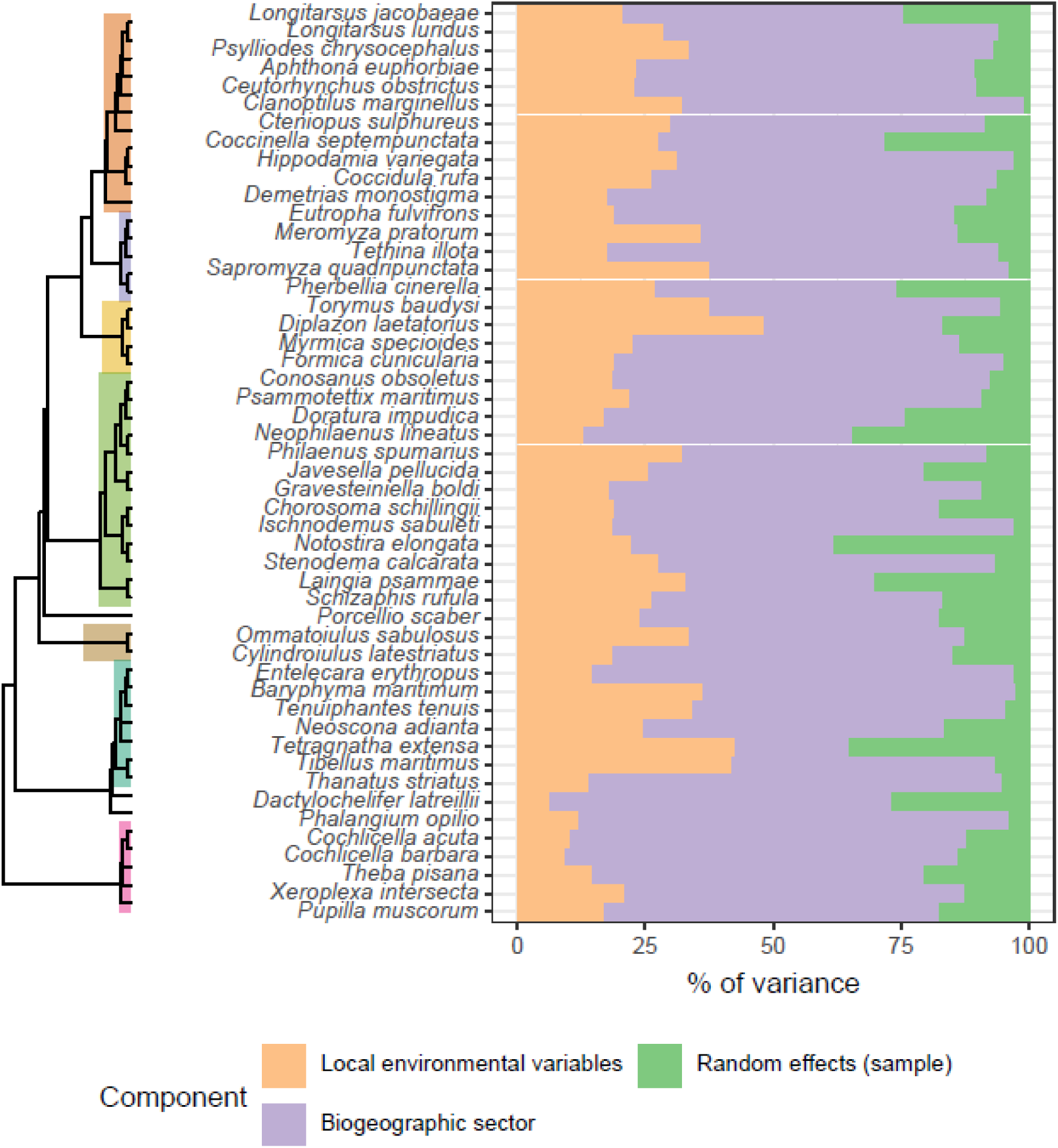
Partitioning of the variation explained by the JSDM. All local variables are pooled for reasons of readability. Key clades are highlighted in the phylogenetic tree using the same colour code as in Figure 2.

### Responses to environmental variables

There were consistent effects of biogeographic sector across species (Fig. 5). Generally, most species were less likely to be present in Dutch and British samples compared to Flemish dunes. In the French Boulonnais region, snails were as likely, on average, to be present than in Flemish dunes. On the other end, snails were consistently less present in British and Dutch dunes, while arthropods’ responses were more variable.

**Figure 5.**
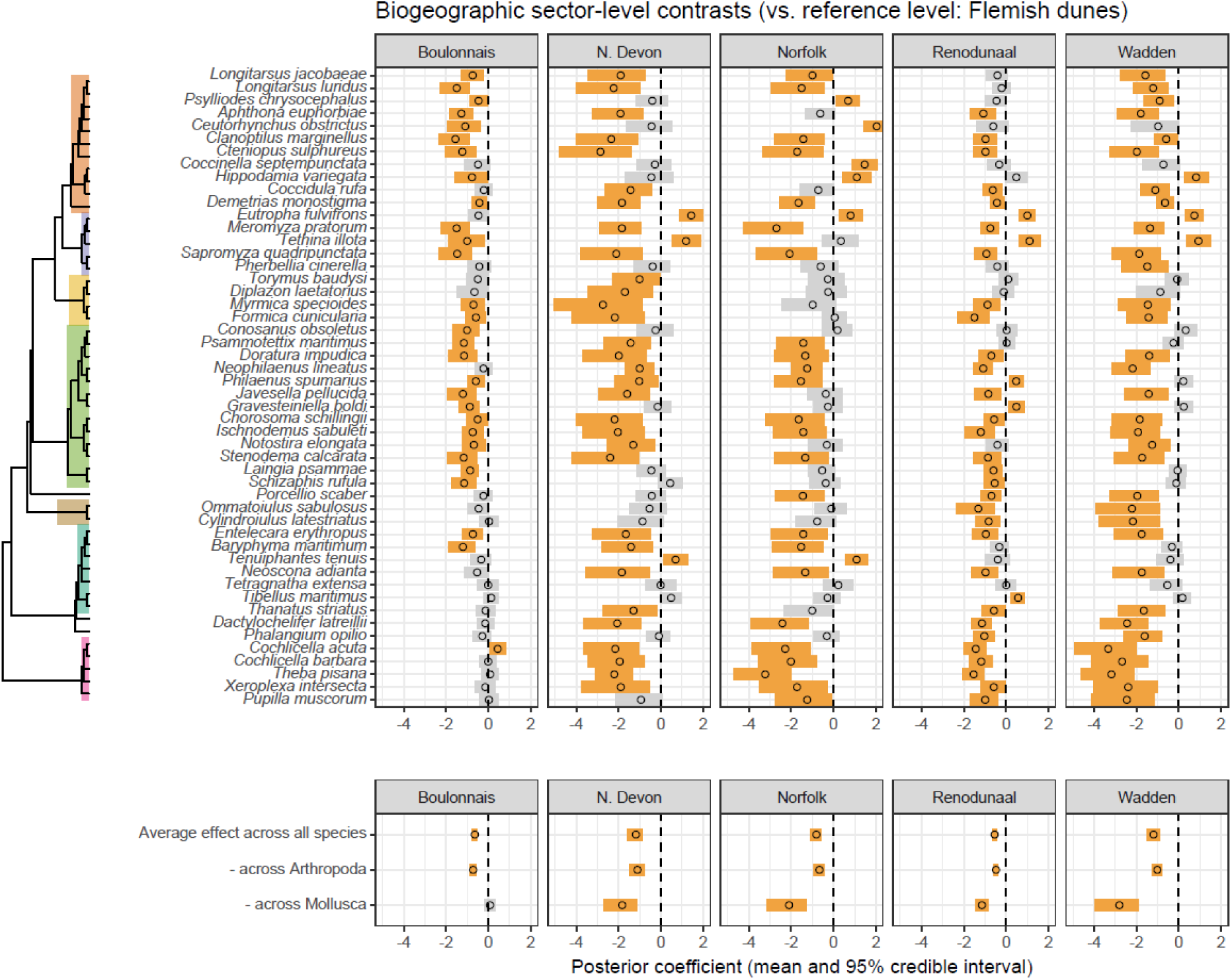
Effects of biogeographic sectors on species probability of occurrence. Posterior means and 95% credible intervals of standardized model coefficients (on the probit scale) are displayed both per species and after averaging across species and by phylum. 95% intervals that overlap with 0 are in grey, intervals that differ from 0 in orange. Key clades are highlighted in the phylogenetic tree using the same colour code as in Figure 2.

Species responses to local environmental variables were more variable (Fig. 6), and for any given niche parameter, the effect on most species was often not different from zero. This was especially true for Moran’s *I*, even though a few species showed negative or positive responses. By contrast, the average linear effect of marram grass cover was different from 0 and positive, and the average quadratic effect negative, indicating potential bell-shaped patterns (Fig. 6, bottom). Effects of marram grass vitality differed between phyla: molluscs showed much stronger responses to vitality, being more likely to occur on lower vitality tussocks. We note that the magnitude of these local effects (based on standardised coefficient values) remains lower than the magnitude of the biogeographical effects (compare x-axis scales on Fig. 5 *vs* Fig. 6).

**Figure 6.**
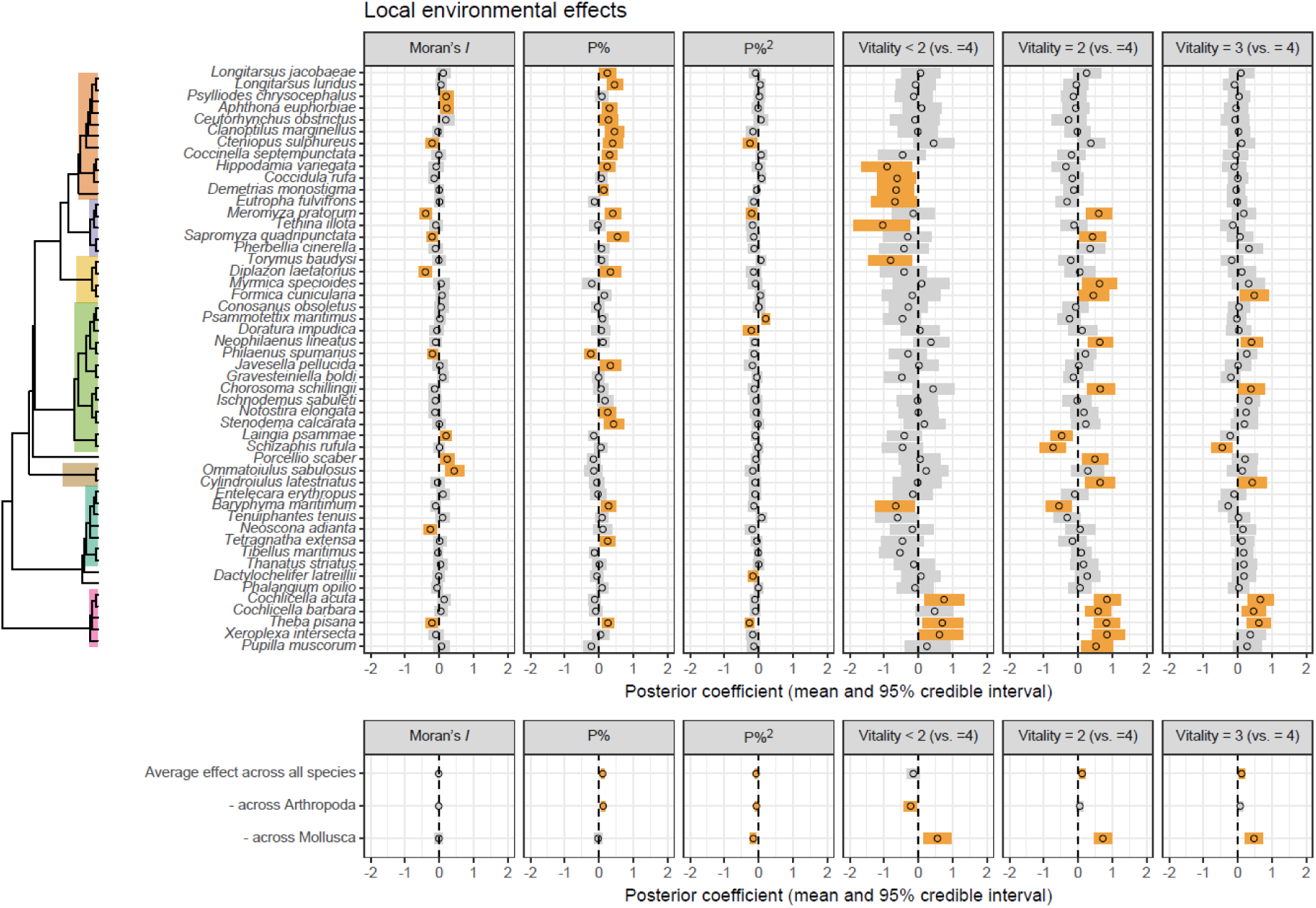
Effects of local environmental parameters (Moran’s I of the marram cover, the percentage cover and its quadratic term P%, P%² and different vitality categories) on species probability of occurrence. Posterior means and 95% credible intervals of standardized model coefficients (on the probit scale) are displayed both per species and after averaging across species and by phylum. 95% intervals that overlap with 0 are in grey, intervals that differ from 0 in orange. Key clades are highlighted in the phylogenetic tree using the same colour code as in Figure 2.

### Species traits and phylogenetic signal

Body size and species functional group generally did not affect species niche in a clear way (Figs. 7-8). One exception was a possible interaction between body size and responses to marram vitality, with larger species more likely to occur on intermediate-vitality tussocks than on high-vitality tussocks (Fig. 8). There was, however, a substantial phylogenetic signal (*ρ* = 0.49 [0.25, 0.68]), meaning that closely related species had more similar responses to the environment than unrelated species (as can be seen Figs. 5-6), which may reflect shared but unmeasured traits.

**Figure 7.**
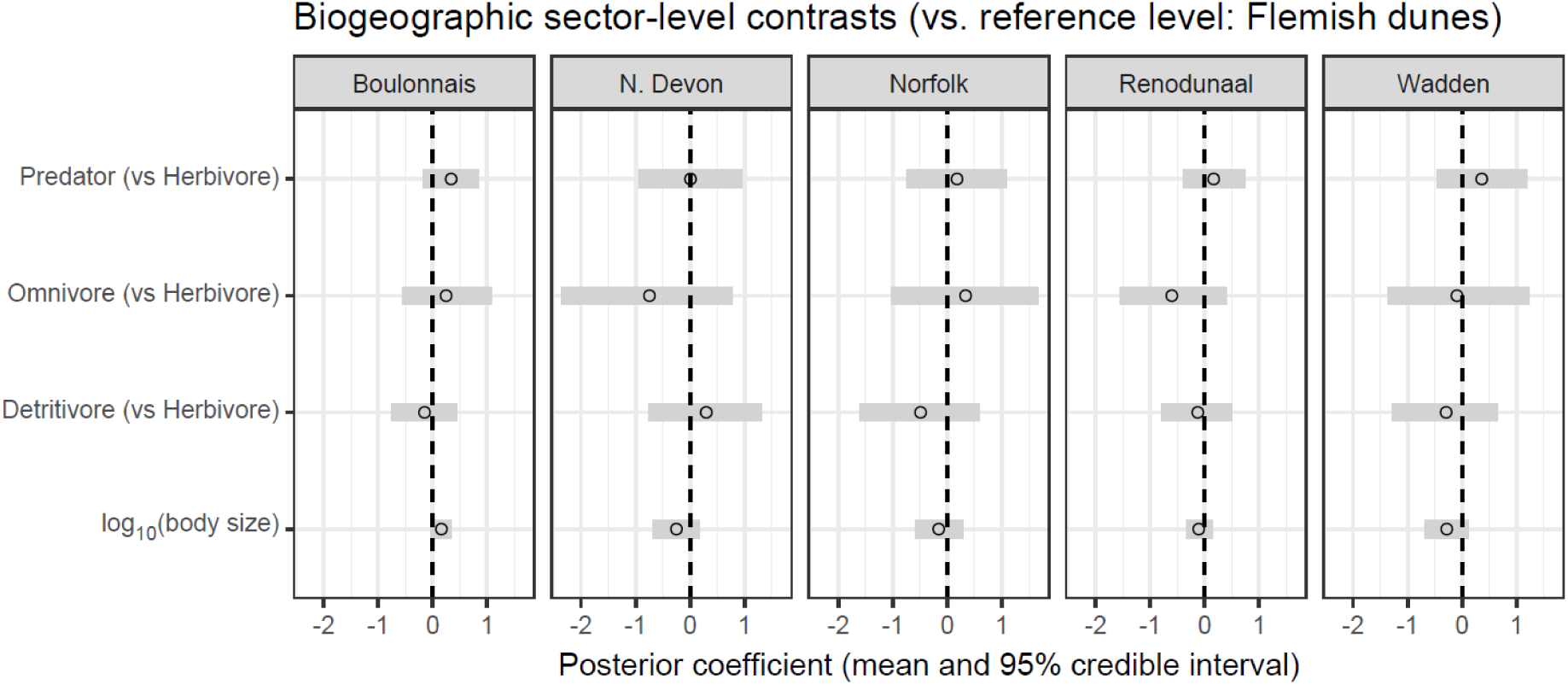
Effect of species traits on species responses to biogeographic sectors. Posterior means and 95% credible intervals of standardized model coefficients (on the probit scale) are displayed.

**Figure 8.**
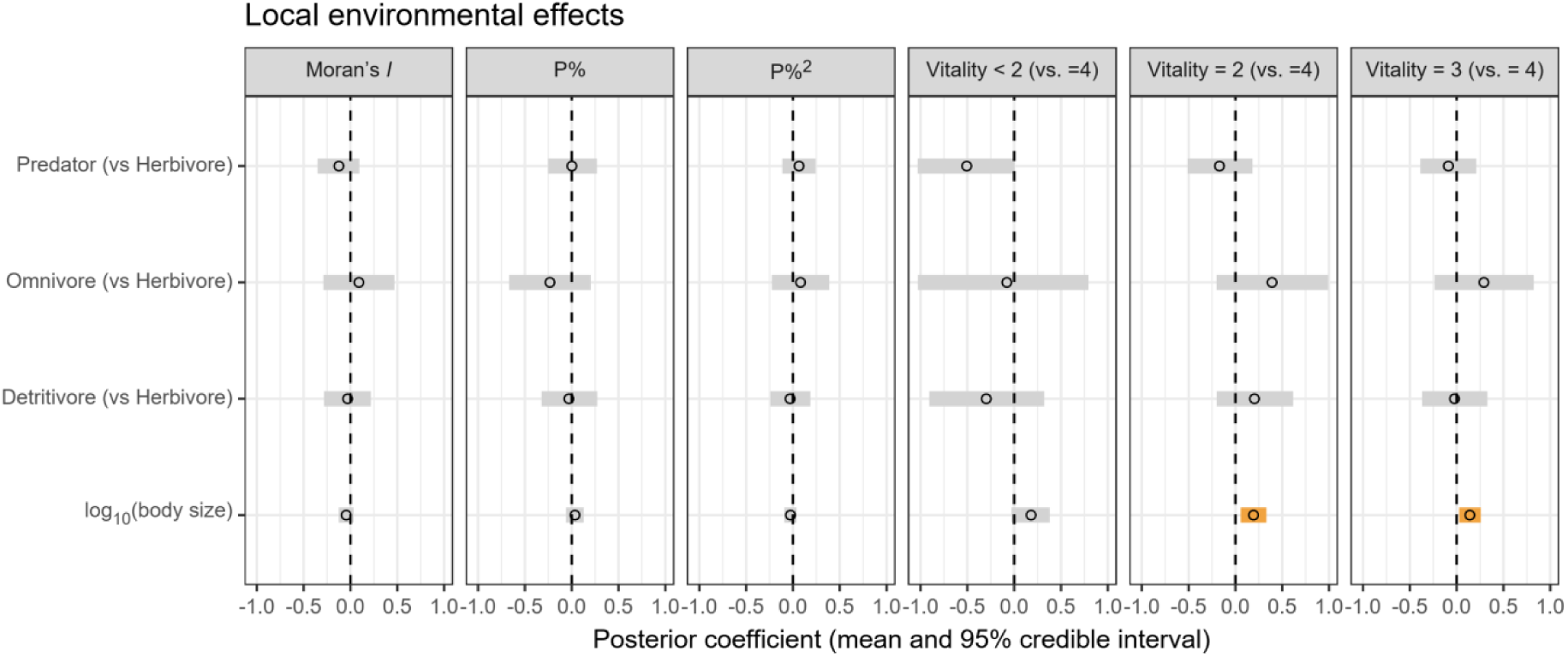
Effect of species traits on species responses to local environmental parameters. Posterior means and 95% credible intervals of standardized model coefficients (on the probit scale) are displayed; 95% intervals that overlap with 0 are in grey, intervals that differ from 0 in orange.

### Residual species co-occurrence patterns

Only strong “residual” correlations (absolute value of the mean posterior > 0.75) were markedly different from 0 (Appendix 5). The correlation matrix showed a clear structure with two separate groups: after accounting for the effect of environmental parameters included in the model, species within one group were positively associated with each other and negatively associated with species in the other group (Fig. 9). The first cluster of co-occurring species contained most of all species, including three of the five most frequent species (*Theba pisana*, *Neophilaenus lineatus*, *Meromyza pratorum*). The second cluster was smaller and included *Schizaphis rufula*, *Eutropha fulvifrons*, *Laingia psammae*, *Ommatoiulus sabulosus*, *Psammotettix maritimus* and *Demetrias monostigma*. The remaining species not belonging to these clusters showed no clear co-occurrence pattern (95% credible intervals for all correlations overlapped 0). Examination of spatial correlation patterns of the site loadings shows evidence of small-scale (< 5 km) autocorrelation in the first latent factor, and none in the other three factors (Appendix 6).

**Figure 9.**
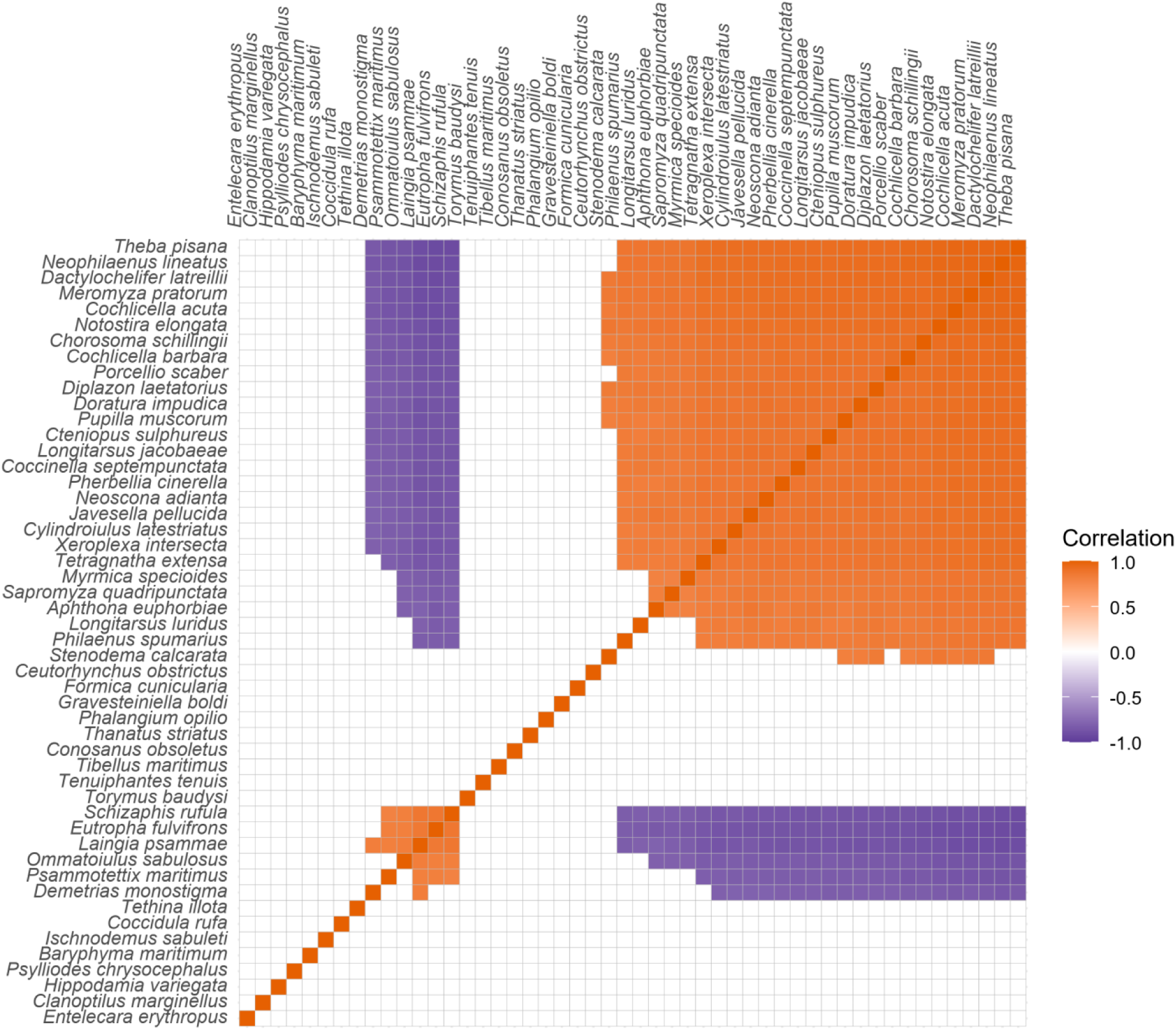
“Residual” species associations, based on sampling unit-level correlations. Species are grouped together based on hierarchical clustering of the posterior mean correlation values (using the average agglomeration method in Hmsc, but different algorithms give similar results). Correlations are only shown if the 95% credible interval did not overlap 0.

## Discussion

Traditional multivariate approaches in community ecology are grounded on distance concepts of the full species assemblage, rather than embracing the complexity of species co-occurrence patterns (Ovaskainen et al. 2017). Such distance-based methods can potentially lead to oversimplified conclusions regarding the environmental drivers of species distributions (Clark et al. 2014, Thorson et al. 2016). Here, we applied both traditional distance-based approaches and a hierarchical modelling framework to understand the drivers of plant-associated invertebrate community structure along a biogeographical gradient along the two-seas region of France, Belgium, the Netherlands and UK. The developed JSDM explicitly models the occurrence of multiple species simultaneously, and sheds light on potential internal drivers of interspecific interactions. Given the many rare species in the system, we unfortunately needed to restrict our analyses to species with a sufficient presence in the full sample set. Despite our extensive stratified sampling, covering more than 600 identified taxa from more than 600 samples, and even when focusing our modelling on the 50 most frequent species, predictions of the occurrence of some -not necessarily the rarest- species are still not significantly better than random patterns, and only very strong “residual” correlations were detected. The used distance-based approaches could handle all abundance data (including the zero inflation) from a minimum total abundance of 15 upward. The RDA analysis identified the same important drivers as the JSDM but remained difficult to interpret because of the nested sampling design. The JSDM’s focus on individual species, their covariation and putative non-linear responses to the environmental gradients enabled, however, a more in-depth variance partitioning among the different factors. PERMANOVA demonstrated the overall importance of the transect as an explanatory factor, and further highlights the importance of local factors related to specific sampled locations as a driver of invertebrate community structure. While this transect information hints at spatial correlation at scales up to 1 km (the transect lengths), it remains speculative to pinpoint exact causes. Given the variation explained by marram grass vitality, cover and spatial clustering, other drivers acting at the transect scale, rather than tussock scale, are in play. Besides inherent stochasticity at this scale, factors related to current and historical management, fragmentation, and recreational use could be of importance.

Besides this local spatial correlation, we show biogeographic sectors to be the most important structuring factor of marram dune invertebrate community structure. JSDM sheds light on the divergent responses of the different species and enables the inclusion of non-linear responses, phylogeny and traits as explanatory factors, while considering correlations in species occurrence. Such insights provide us more accurate and comprehensive predictions of species distributions in light of e.g. nature restoration schemes and biodiversity monitoring by remote sensing of vegetation patterns (Cavender-Bares et al. 2022).

This species-specificity could in general not be explained by species body size, although a phylogenetic signal remained prominent. Related species thus showed similar responses in relation to the environmental variables, indicating non-random associations between dune invertebrates and the cover and vitality of the host marram grass plants. This non-random association is also reflected in the residual species associations, indicating two clusters of positive and negative associations. The retrieved spatial correlation indicates here as well that if the species association patterns are due to missing environmental variables, the key ones are probably varying at scales varying from local (within transects) up to a few kilometers (neighbouring transects).

The identified biogeographic sectors differ in their geological and geographical structure as reflected in the sediment chemical composition, especially lime richness and thus soil pH, the prevalence of a well-documented north-south gradient of climatic variables (e.g., temperature) and UK-mainland separation (Bonte et al. 2003a). Such environmental variation also impacted the historical use of these systems in different ways (Provoost & Bonte 2004). These factors are known to drive species community structure of both plants (e.g., Forey et al. 2008) and animals (e.g., Bonte et al. 2003a) from grey dunes, also being stressed coastal dune habitats. We show here that large spatial heterogeneity is also a driver for invertebrate communities from systems (e.g., blond dunes) that experience strong natural disturbance by aeolian dynamics. It is not so speculative to assign the strong association of snails with the Flemish and the Boulonnais dunes to lime availability (Graveland and Wal 1996, Provoost and Bonte 2004). The occurrence of the generalist spider species *Tenuiphantes tenuis* in marram dunes could only be linked to the distinction between the European continent and the UK samples, pointing at either niche differences between the two populations or specific conditions allowing their persistence in the UK dunes, while being only rare vagrants in coastal dunes of the continent. Since this species is a strong disperser by air (ballooning; Bonte et al. 2003b, 2004), prevailing landward winds typically prevent the species from drifting into the coastal systems. However, in the UK, the pattern of mass immigration via wind-driven drift may be more pronounced. We need to note that different countries (and by extension, biogeographic sectors) were sampled during different years, implying that such a reasoning remains very speculative as it may be equally caused by weather conditions prior to the sampling (Didham et al. 2020). Also, compared to mainland sand dunes, UK dunes were relatively undersampled because of the overall rarity of developed coastal dunes in the region. Rerunning the final model with only data gathered during 2018, or without UK samples, confirmed the biogeographic sector as a robust explanatory factor.

90 % of the species showed associations with spatial configuration of the marram vegetation as measured by its cover (P%), spatial clustering (Moran’s *I*) and/or the marram grass vitality. Such important insights cannot be obtained from the full distance-based community approaches. Responses of all other species were heterogeneous, indicating species-specific sorting mechanisms among niches associated with these recorded environmental variables. These environmental factors are affected by aeolian dynamics (Bonte et al. 2021), which in turn have been demonstrated to be a strong driver of species composition along dune successional gradients (Bonte et al. 2006, Forey et al. 2008). In dynamic blond dunes such as those studied here, marram grass tussocks are highly clustered due to self-organising mechanisms (Bonte et al. 2021). Variability in spatial clustering is therefore naturally low, and hence also the explained variation relative to the other recorded environmental factors. Responses to marram grass cover (P%) were always positive or parabolic, indicating that chances of detecting species increase with cover (and thus availability of plant-associated resources), or show an optimum value. The positive linear responses indicate that marram grass tussocks function as islands in a hostile matrix according to the theory of island biogeography (TIB), with larger islands having higher species richness (MacArthur and Wilson 1967, Gravel et al. 2011). Most species showing such responses can be regarded as being generalist and only loosely associated with marram grass as their host. Species with a hump shaped response are already more dune- and sand-associated (e.g., *Theba pisana, Doratura impudica*), indicating their need for a sufficiently heterogeneous sand-vegetation environment. Only one species was preferentially associated with a low marram grass cover; the species *Philaenus spumarius* does not have an affinity for open sand, so the exact reason of this association remains very speculative. Plants in low cover may be stressed and therefore less resistant against such herbivory, but then other invertebrate species should be expected to show the same pattern. Alternatively, the low cover may provide the thermal properties preferred by this species. Associations with marram grass vitality is clearly more phylogenetically structured and reflects trophic positions with more grass-bound sap suckers (aphids *Laingia psammae and Schizaphis rufula*) and their predators (*Demetrias monostigma* and *Baryphyma maritimum*; Van De Walle et al. 2023) associated with the most vital marram grass tussocks. Conversely, more widespread detritivores like the isopod *Porcellio scaber* and some snail species (*Cochlicella* spp. and *Pupilla muscorum*) preferred marram grass tussocks with more detritus as they use dead plant material as hiding place, food source or a combination of both. We re-ran phylum-specific models to determine whether the observed phylogenetic signal was primarily due to differences between Arthropods and Molluscs. The latter show indeed a distinct underrepresentation in some (less calcareous) biogeographic sector (Fig. 6). These models did not fully converge despite increasing iterations compared to the base model, again highlighting the limits of JSDM when data are limited (see Data availability for details). Phylogenetic signal parameters did converge, with values of 0.37 [0,0.65] for arthropods and 0.46 [0, 0.99] for molluscs, which does not allow us to conclude whether between-species differences in niches are primarily due to differences between Arthropods and Molluscs.

The use of JSDM has the advantage of retrieving residual covariation among species that may inform on the relevance of species interactions as a driver of community structure (but see Poggiato et al. 2021). We retrieved the existence of two groups of invertebrates that are positively co-occurring within, but negatively between groups. The larger group contained generalist species which are not specifically associated with marram grass or even with dunes or warm habitats in general, such as *Coccinella septempunctata* (a ladybird), *Longitarsus jacobaeae* (a leaf beetle), *Notostira elongata* (a capsid bug), *Philaenus spumarius* (a spittlebug) and *Porcellio scaber* (a woodlouse). The smaller group mainly consisted of dune specialists which strongly depend on marram grass for their survival. For instance, *Laingia psammae* and *Schizaphis rufula* are two aphids associated with grasses and common on marram grass (Weeda et al. 1991, Vandegehuchte et al. 2010), *Psammotettix maritimus* (a leafhopper) feeds exclusively on marram grass (Nickel 2009) and *Eutropha fulvifrons* (a grass fly) uses marram grass as food plant (Nartshuk and Andersson 2013). The remaining species occurred independently from the other groups and from each other. This group seemed to consist largely of predatory (spider) species such as *Tenuiphantes tenuis*, *Thanatus striatus*, *Entelecara erythropus* and *Tibellus maritimus*. Since the effect of marram grass tussock vitality is already considered, this result could point towards priority effects between specialist and generalist species rather than succession. This would entail that community assembly depends on the order and timing in which species form and join communities (Chase 2003, Fukami 2015). This process is plant-mediated, with the hostplant becoming more suitable for either of the two communities, depending on the first species to arrive. Marram grass might, for instance, allocate more resources to the roots as a response to above- ground herbivory (Kaplan et al. 2008), or induce (primary) defence across the above- and belowground compartments (Vandegehuchte et al. 2010). With this explanation, it is perfectly understandable that the remaining species not co-occurring with either group are mainly predators.

Alternatively, this result could be explained by the effect of an environmental or genetic variable not included in the model. If relevant, this missed variable should be expressed at relatively small spatial scales up to 5 km (Appendix 6), just as retrieved for the transect- variation in the distance-based analysis. This scale excludes e.g. environmental variables such as invasive plant distributions (Van De Walle et al. 2022). Most of the coastal dunes have experienced episodes of marram grass planting, potentially obscuring natural genetic variation linked to above-belowground interactions (Vandegehuchte et al. 2011) at smaller spatial scales. Alternatively, unmeasured variation in microclimate or belowground biotic interactions (van der Putten et al. 2009) can induce different plant-mediated defence responses that potentially induce the observed ecological sorting between specialist and generalist feeders.

Marram grass spatial clustering, cover and vitality are directly linked to the prevailing sand dynamics (Huiskens 1979, Nolet et al. 2018, Reijers et al. 2021, Bonte et al. 2021, Strypsteen et al. 2024) and these days directly managed through planting actions to stabilise existing dunes and/or promote the evolution of new dunes in front of urban areas as a Nature-based Solution for coastal protection (van der Meulen et al. 2023). These actions consequently have an immediate impact on the system’s biodiversity value. Blond dunes with an intermediate cover and clumped distribution of marram grass have been demonstrated to maximise dune growth (Bonte et al. 2021, Strypsteen et al. 2024). Hence, it seems that based on both biodiversity values and putative coastal protection services, unvegetated nourishment at the upper beach should be cautioned against, as barely vegetated dunes sustain neither biodiversity nor dune development. Rather, total arthropod species diversity is maximised when dunes are completely vegetated, so in grass-encroached dunes, while ecosystem functioning with respect to dune growth is maximised when patches of bare sand and vegetation alternate. This apparent conflict arises from invertebrate species richness increasing under succession, while aeolian dynamics and sand deposition are maximised when marram grass cover reaches approximately 50%. Species of conservation concern, narrowly associated with coastal dunes show either hump-shaped responses to marram cover, or a positive response to marram vitality. Because marram vitality is maximal under sand burial (Bonte et al. 2021), we can state that optimal states for dune development and resilience align with the conditions needed for species of conservation concern. Hence, rather than a conflict, our results thus suggest that optimal states for the restoration of coastal protection services and dune-specific biodiversity converge.

As a final point of attention, both approaches informed us on the importance of local ‘transect’ factors as important drivers of invertebrate community similarity. Hence, besides the small-scale individual-plant variables and the large-scale biogeography, some intermediate-scale drivers are clearly at play. These should operate at the metacommunity level and are most likely related to spatial drivers limiting species spread within the fragmented dune area in which the transects were located, in combination with local factors related to dune management and/or recreational use (e.g., Bonte et al. 2003a, 2004, Maes and Bonte 2006, Bonte and Maes 2008). Such human impacts may even occur from a legacy of past management. For instance, it has been demonstrated that marram grass genotypes, and the location of origin of planted material, strongly impact above- and belowground plant-associated invertebrates (Vandegehuchte et al. 2011). Planting actions in the past to stabilise foredunes may therefore be one of the most important causes of invertebrate community structure. Unfortunately, no data on such management, let alone the origin of the planted material, or their genetics are currently available.

## Acknowledgements

Part of this research has received funding from the European Interreg 2-seas programme (Endure, No 2S03-007) and European Union’s Horizon Europe research and innovation programme under grant agreement No 101135410 – The DuneFront Project. DB was funded by BOF (grant nr: BOF/24J/2021/066). FM would like to thank CNRS for general support. We thank everyone who has assisted with the fieldwork and/or identification of the arthropods: Fons Verheyde, Paulien Vanhauwere, Laurian Van Maldegem, Gillis Sanctobin, Cyr Mestdagh, Noëmie Van den Bon, Hans Matheve, Pieter Vantieghem and the Endure consortium. We especially would like to thank Lorenzo Munari for assistance in the identification of Canacidae.

## Data availability

Data and R scripts to reproduce all analyses presented in this article are available on GitHub (https://github.com/mdahirel/JSDM_dune_arthropods_vdw) and archived in Zenodo (https://zenodo.org/doi/10.5281/zenodo.12079615).

## Author contributions

MLV and D. Bonte designed the field work. RVDW and MLV conducted the practical work. WL, MD and RVDW wrote the first draft of the manuscript. WL, D. Benoit, D. Bonte, RVDW and MD analysed the data. All authors contributed substantially to interpretation of the results and revision of the manuscript.

## Appendices

## Appendix 1 : Climatological differences among the biogeographic sectors

**Figure S1-1.**
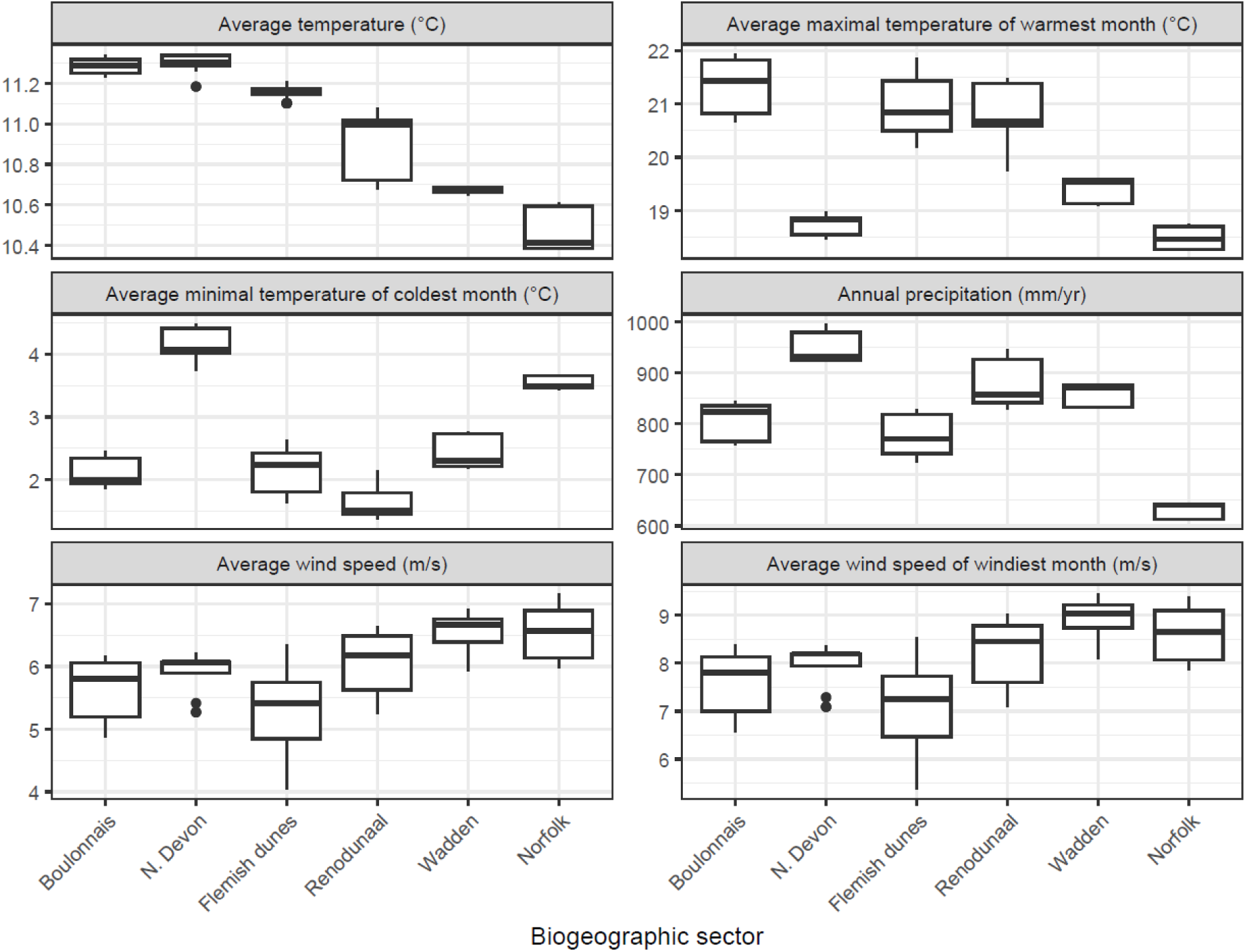
Average climate variables (over the 1989-2018 climatological period) at each sample, grouped by biogeographic sector (data from CHELSA, version 2.1, Karger er al. 2017, 2018). Biogeographic sector are ordered approximately by increasing latitude.

*References:*

*Karger, D.N., Conrad, O., Böhner, J., Kawohl, T., Kreft, H., Soria-Auza, R.W., Zimmermann, N.E., Linder, P., Kessler, M. (2017): Climatologies at high resolution for the Earth land surface areas. **Scientific Data.** 4 170122. https://doi.org/10.1038/sdata.2017.122*

*Karger D.N., Conrad, O., Böhner, J., Kawohl, T., Kreft, H., Soria-Auza, R.W., Zimmermann, N.E, Linder, H.P., Kessler, M. (2018): Data from: Climatologies at high resolution for the earth’s land surface areas. **EnviDat**.https://doi.org/10.16904/envidat.228.v2.1*

## Appendix 2 : References used for species identification & their traits

### General

Provoost, S., & Bonte, D. (2004). Levende duinen, Een overzicht van de biodiversiteit aan de Vlaamse kust.

### Hemiptera

Biedermann, R. & R. Niedringhaus (2004). The plant- and leafhoppers of Germany. Identification key to all species. WABV, Scheeßel, Osnabruck. 409pp.

Kunz, G., H. Nickel & R. Niedringhaus (2011). Fotoatlas der Zikaden Deutschlands - Photographic Atlas of the Planthoppers and Leafhoppers of Germany. WABV Fründ, Osnabruck. 293pp.

Vandegehuchte, M. L., de la Peña, E., & Bonte, D. (2010). Aphids on Ammophila arenaria in Belgium: First reports, phenology and host range expansion. Belgian Journal of Zoology, 140(1), 77–80.

### Heteroptera

Aukema, B., Cuppen, J., Nieser, N., & Tempelman, D. (2002). *Verspreidingsatlas Nederlandse wantsen (Hemiptera: Heteroptera). Deel I: Dipsocoromorpha, Nepomorpha, Gerromorpha & Leptopodomorpha.* European Invertebrate Survey-Nederland.

Southwood, T. R. E. & Leston, D. (1959). Land and water bugs of the British Isles., Frederick Warne & Co. Ltd., London & New York.

### Coleoptera

Bieńkowski, A. O., & Orlova-Bienkowskaja, M. J. (2015). Trophic specialization of leaf beetles (Coleoptera, Chrysomelidae) in the Volga Upland. Biology Bulletin, 42, 863–869. https://doi.org/10.1134/S1062359015100015

Delbol, M. (2010). Les Otiorhynchini de Belgique (Curculionidae : Entiminae). Entomologie Faunistique – Faunistic Entomology, 62(4), 139–152.

Cuppen, J., Kalkman, V., Tacoma, G., & Heijerman, T. (2015). Veldklapper Lieveheersbeestjes. EIS Kenniscentrum Insecten en andere ongewervelden.

Muilwijk, J., Felix, R., Bleich, O. and Dekoninck, W. 2015. De Loopkevers van Nederland en België (Carabidae). - Nederlandse Entomologische Vereniging, Naturalis Biodiversity Center.

Schmitt, M. (1988). The Criocerinae: Biology, Phylogeny and Evolution. In P. Jolivet, E. Petitpierre, & T. H. Hsiao (Eds.), Biology of Chrysomelidae. Springer Netherlands. https://doi.org/10.1007/978-94-009-3105-3

### Diptera

Munari L. (2011). The euro-mediterranean Canacidae s.l. (including Tethinidae): keys and remarks to genera and species (Insecta, Diptera). Boll Mus. St. Nat. Venezia, 62: 55-86.

Nartshuk, E. P. and Andersson, H. (2013). The Frit Flies (Chloropidae, Diptera) of Fennoscandia and Denmark. Fauna Entomologica Scandinavica, 43, Brill Academic Publishers, Leiden. 277pp

Oosterbroek P. (2006). The European families of the Diptera. Identification, diagnosis, biology. KNNV Publishing, Utrecht, 205 pp.

Rozkošný, R. (1984). The Sciomyzidae (Diptera) of Fennoscandia and Denmark. Fauna Entomologica Scandinavica 14, Scandinavian science press Ltd, Klampenborg. 224pp.Schacht, W., Kurina, O., Merz, B., & Gaimari, S. D. (2004). Zweiflugler aus Bayern XXII (Diptera: Lauxaniidae, Chamaemyiidae).

Entomofauna Zeitschrift Fur Entomologie, 25, 41-80.

Shatalkin, A. I. (2000). Keys to the Palaearctic flies of the family Lauxaniidae (Diptera). Zoologicheskie Issledovania, 5, 102pp.

### Other families

Ellis, W. N. (2001). Bladmineerders. Bladmineerders.nl

Crumb, S. E., Eide, P. M., & Bonn, A. E. (1941). The European Earwig. Technical Bulletin.

Roberts, M. (1985). The Spiders of Great Britain and Ireland Compact Edition (2 vols.). Brill publishing

## Appendix 3 : MCMC chain diagnostics

**Figure S3-1.**
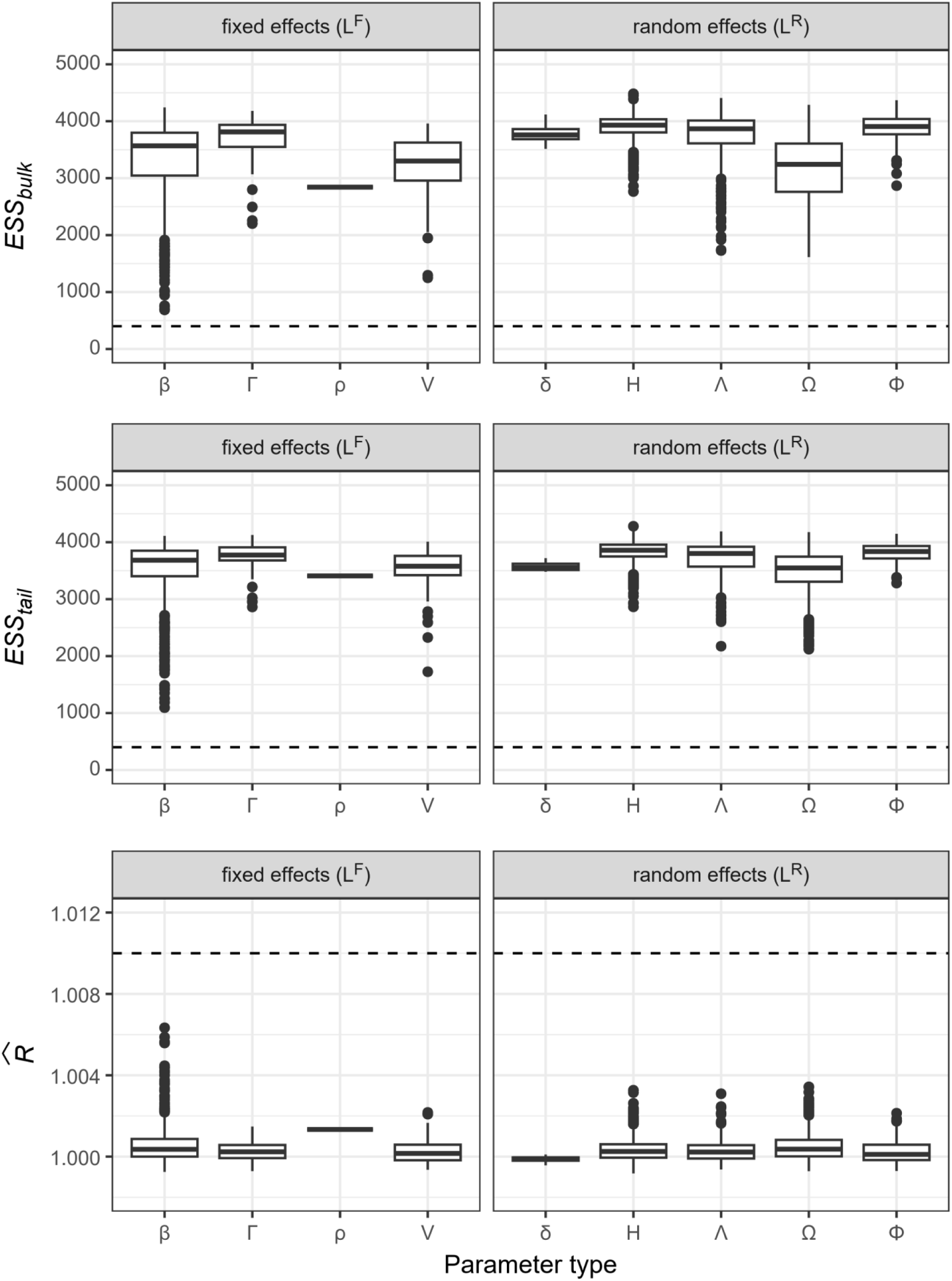
Effective sample sizes (bulk and tail) and potential scale reduction factor *R̂* (sensu Vehtari et al. 2021) for all model parameters. Dashed lines are the thresholds above (effective sample size) or below (*R̂*) which parameters can be deemed satisfactory. Parameter groupings and names are as in Ovaskainen and Abrego (2020).

## Appendix 4 : Model performance

**Figure S4-1.**
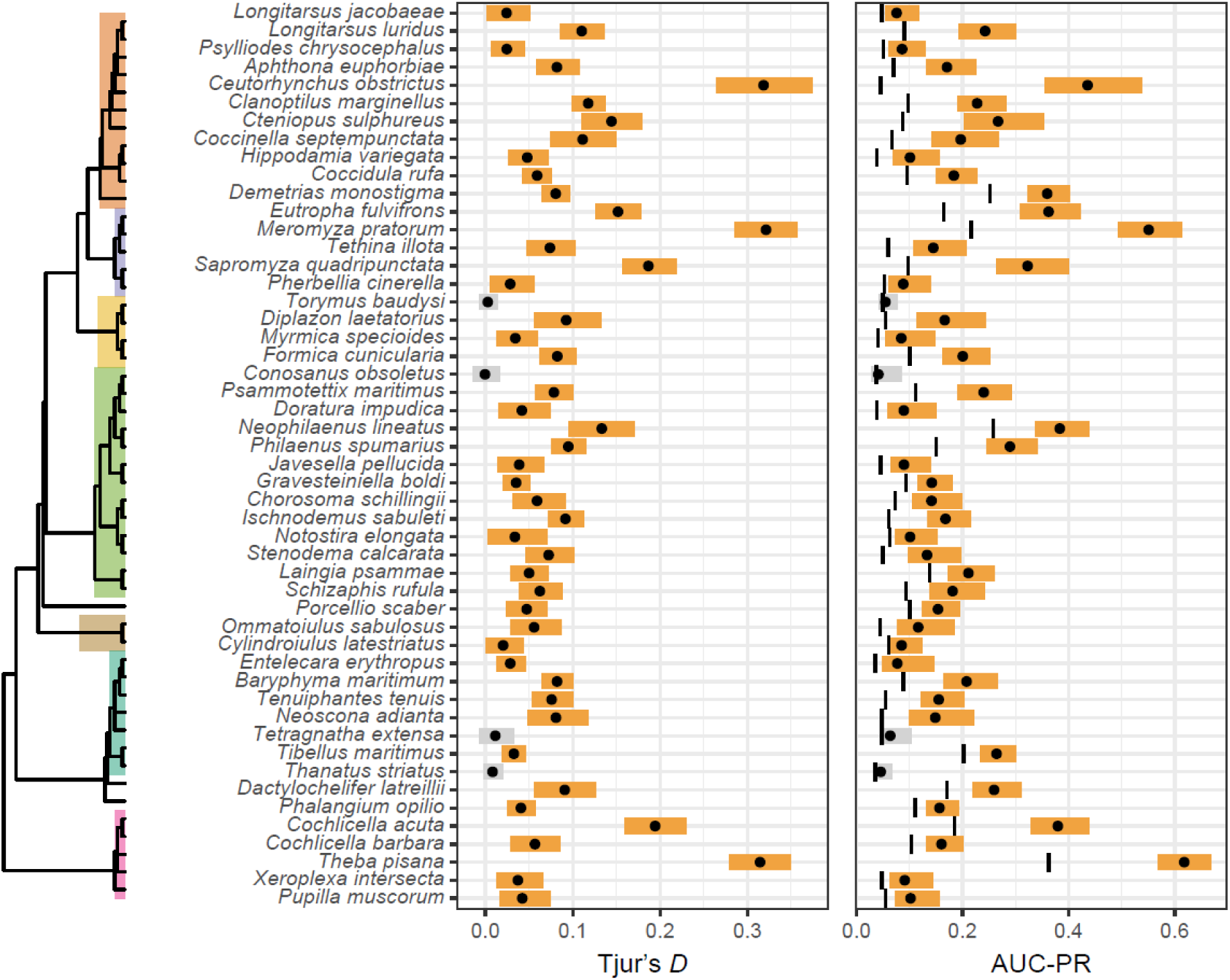
Distribution of Tjur’s D and AUC-PR values among species, based on predictions made after 10-fold cross-validation. The vertical segments on the AUC-PR panel denote the observed prevalence, which is the expected value if the model performs at random. Coloured bands are 95% credible intervals per species; 95% intervals that overlap with the random expectation are in grey. Key clades are highlighted in the phylogenetic tree as in main text Figure 2.

**Figure S4-2.**
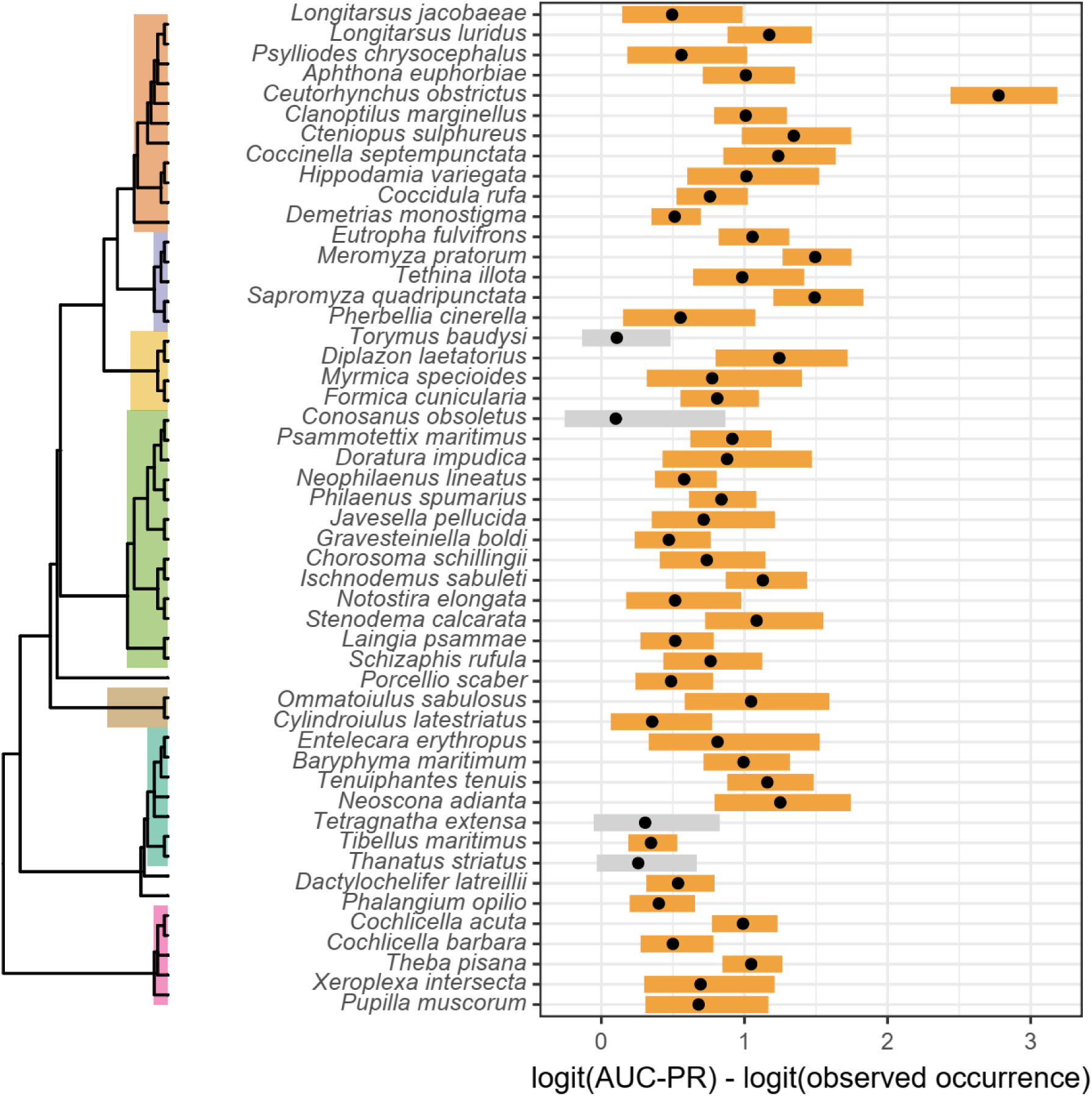
Difference on the logit scale between out-of-sample model performance for each species (measured by cross-validated AUC-PR) and its expected values for a model performing at random (equal to observed occurrence). Key clades are highlighted in the phylogenetic tree as in main text Figure 2.

## Appendix 5 : threshold to detect “residual” species correlations

**Figure S5-1.**
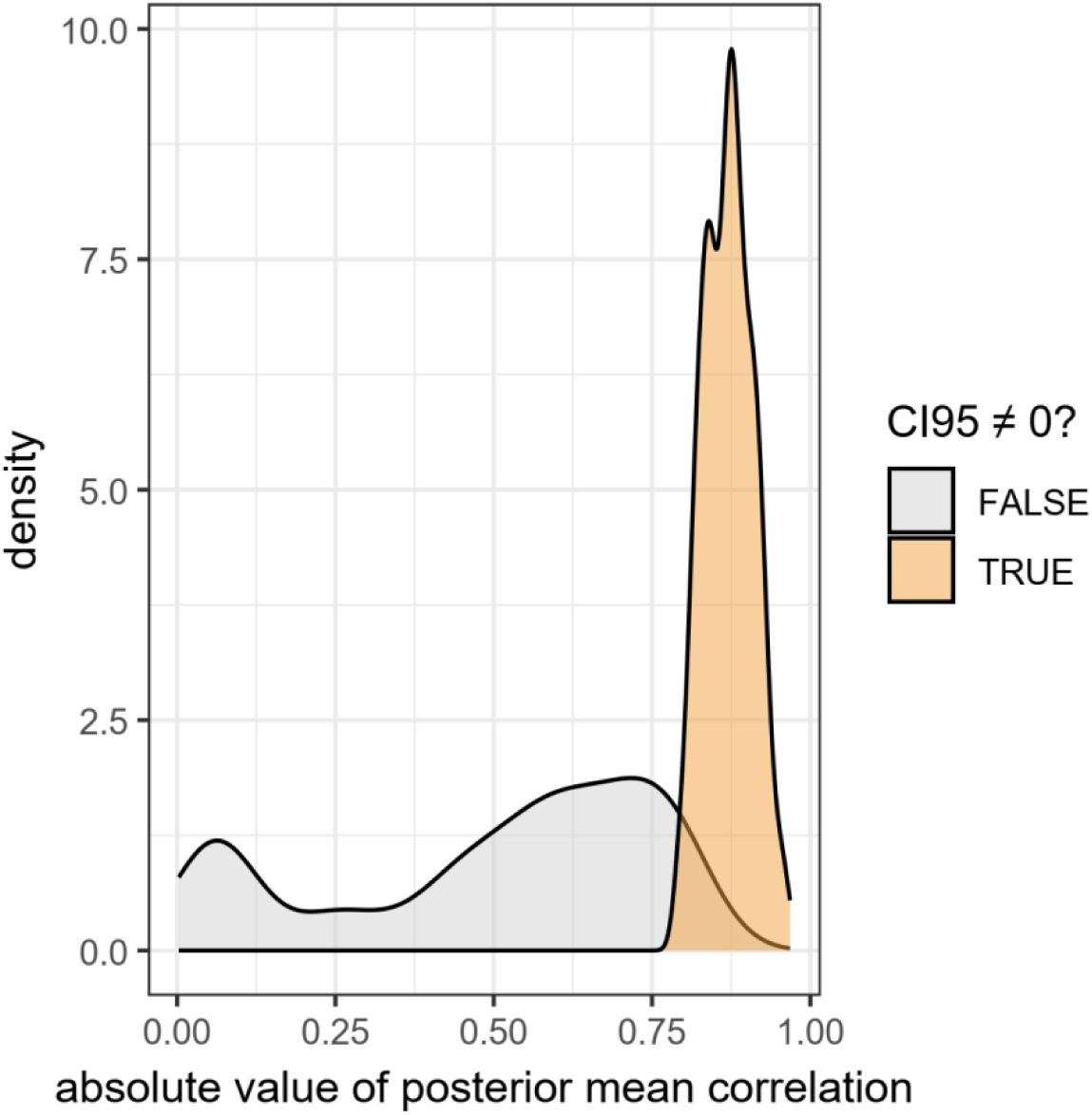
Distribution of absolute mean posterior values of species-species correlations for which the 95% credible interval did and did not overlap 0. Only strong correlations are markedly different from 0.

## Appendix 6 : spatial autocorrelation in site loadings

**Figure S6-1.**
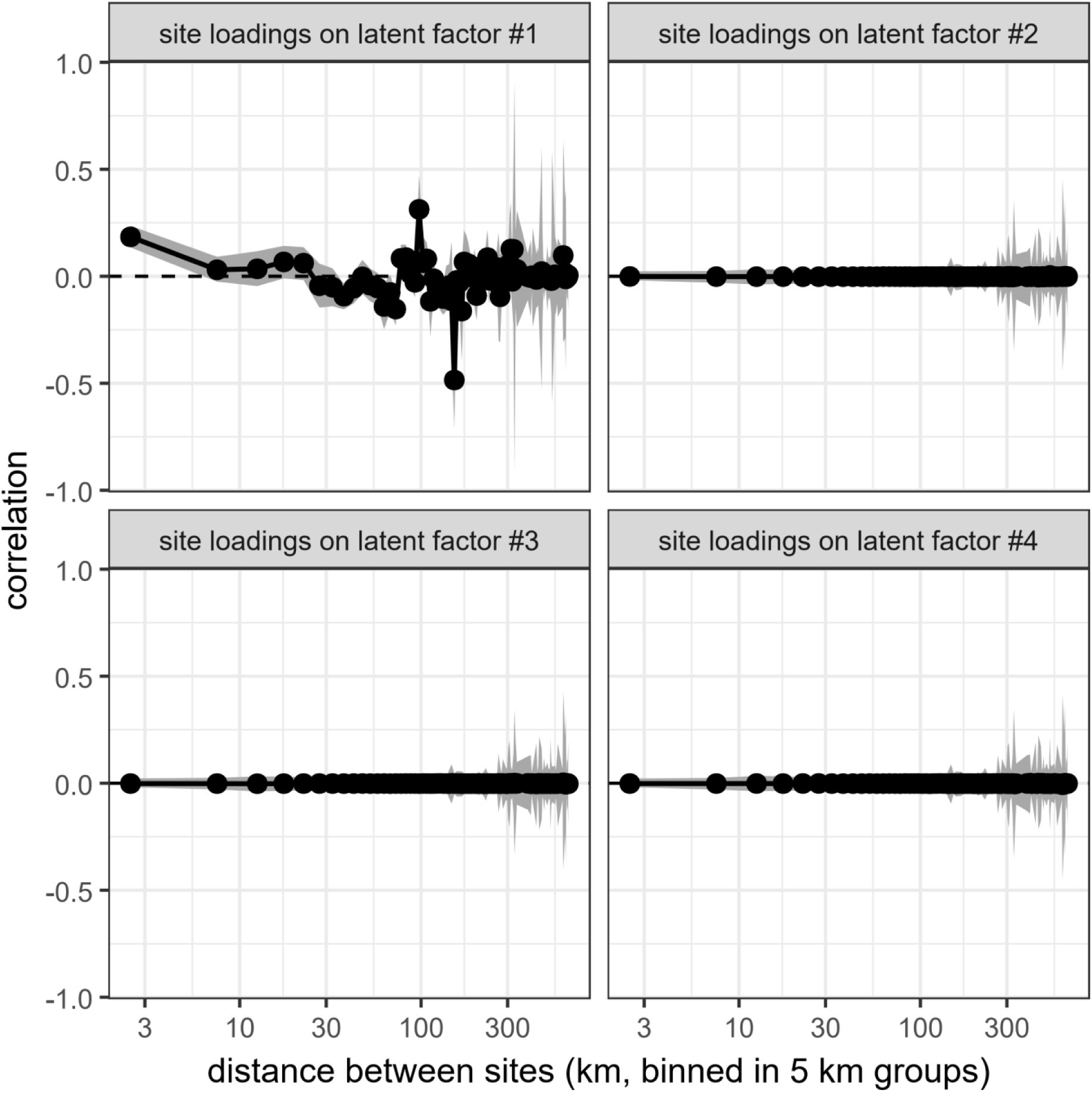
Spatial correlograms showing the correlation in site loadings (H in Appendix 2) as a function of distance between sites. Spatial dependence values are estimated for each posterior sample, the means and 95% credible intervals for each distance class are displayed. Note the log10 scale on the x axis.

